# Nafamostat Mesylate in lipid carrier for nasal SARS-CoV2 titer reduction in a hamster model

**DOI:** 10.1101/2020.11.09.372375

**Authors:** Lisette Cornelissen, Esmee Hoefsmit, Disha Rao, Judith Lijnsvelt, Lucien van Keulen, Marieke van Es, Volker Grimm, René H. Medema, Christian U. Blank

## Abstract

Severe acute respiratory syndrome corona virus 2 (SARS-CoV-2) has been responsible for the largest pandemic in recent decades. After seemingly being in control due to consequent lock-downs and social distancing, the majority of countries faces currently a second wave of exponentially increasing infections, hospital referrals and deaths due to SARS-CoV-2-mediated disease (COVID-19). To date, no effective vaccination has been found, and wearing masks and social distancing are the only effective approaches to reduce further spreading.

However, unwillingness in the societies to distance again and consequently wear masks might be reasons for the second SARS-CoV-2 infection wave. User-friendly chemicals interfering at the host site with viral entry might be an approach to contain the pandemic. In addition, such an approach would work synergistic with vaccinations that miss new virus mutants.

Nafamostat (NM) has been shown *in vitro* to interfere with cellular virus entry by inhibition of the host transmembrane protease serine 2 (TMPRSS2), an enzyme required for SARS-CoV-2 spike protein cleavage, a prerequisite for cell entry.

We hypothesized that nasal application of NM in a liposomal layer (as additional mechanical barrier) could lower the nasal viral load and subsequently reduce the severity of COVID-19. We found, indeed, that nasal viral load one day post single NM application, was lowered in a hamster SARS-CoV-2 infection model. However, severity of subsequent local tissue destruction and weight loss due to pneumonitis was not favorably altered.

In conclusion, a single NM application reduced nasal viral load, but did not favorably improve the outcome of COVID-19, likely due to the short half-time of NM. Improvement of NM stability or repetitive application (which was not permitted in this animal model according to Dutch law) might circumvent these challenges.

## Introduction

At the end of 2019 a novel coronavirus, severe acute respiratory syndrome corona virus 2 (SARS-CoV-2) has been identified for the first time in Wuhan, China, being the causative agent of the ongoing pandemic Coronavirus Disease 2019 (COVID-19). This virus has been responsible for the largest pandemic in modern times with more than 42 million infections demanding more than a million deaths (John Hopkins Coronavirus Resource Center, 10/2020). SARS-Cov-2 is thought to be predominantly transmitted in nanoparticles after close contact with an infected person.

The spike (S) protein of coronaviruses facilitates viral entry into target cells. Entry depends on binding of the surface unit S1 of the S protein, to a cellular receptor, which facilitates viral attachment. SARS-S has been shown that the angiotensin-converting enzyme 2 (ACE2) is the entry receptor (*1*), which has been proven to be also the case for SARS-CoV2 (*2*). High-sensitivity RNA in-situ mapping revealed the highest ACE2 expression in the nose with decreasing expression throughout the lower respiratory tract, paralleled by a striking gradient of SARS-CoV-2 infection in proximal (high) vs distal (low) pulmonary epithelial cultures (*3*). This expression pattern explains the first symptoms of infections often being congestion or runny nose, new loss of taste or smell, and sore throat.

In addition, entry requires S protein priming by host cellular proteases, which entails S protein cleavage at the S1/S2 and the S2’ site and allows fusion of viral and cellular membranes, a process driven by the S2 subunit. SARS-Cov-2 employs the cellular serine protease TMPRSS2 for S protein priming (*2*).

Currently, several groups are working on developing vaccines against SARS-CoV-2, often targeting S protein subunits. These vaccines are expected at earliest next year to be available to broader populations. Re-infections after COVID-19 are to date rare and have been described in only 3 patients (*4–6*). In two of them, phylogenetically distinct SARS-CoV-2 viruses were isolated, both harboring mutations in the spike protein, possibly questioning life-long immunization from the current vaccination strategies. This worry is further supported by recent COVID-19 infections in mink in Denmark that carry mutated strains that might be able to spread from mink to humans (*7*).

Meanwhile social distancing, wearing masks, washing hands regularly, intense testing for infections, and quarantining infected persons are the most frequent advices from virologists and epidemiologists and are generally accepted approaches in most countries of the world. However, after the first lock-down beginning 2019, accompanied with reduced death rates in almost all countries, a second wave is observed in the recent months after lowering the contact restrictions (ig.ft.com/coronavirus-chart). In many countries, demonstrations against the strict rules are occurring, indicating the disappearing willingness to restrict to social distancing and wearing masks. In addition, incorrect use of face masks might also fail to prevent virus transmission.

Possibly interim solutions (until a broad vaccination initiative is available) might be approaches inhibiting entry of the virus in the upper respiratory tract. Nafamostat Mesylate (NM) has been developed and approved as anticoagulant in Japan (*8*). NM has been identified a potent inhibitor of coronavirus S protein mediated membrane fusion (*9*), and has been shown *in vitro* to inhibit SARS-CoV-2 entry into cultured human lung epithelial cells by inhibiting TMPRSS2 activity (*10,11*). In addition, such chemical virus entry inhibition targeting host enzymes could work synergistically with vaccination, local nanobody delivery (*12*), or inhibitory lipopeptides (de Vries et al., bioRxiv preprint doi: https://doi.org/10.1101/2020.11.04.361154), and still be effective in case of SARS-CoV-2 escape variants occur from vaccination.

Liposomes layers applied by nose sprays have been shown to reduce seasonal-allergic-rhinoconjunctivitis (*13*). We hypothesized that application of NM, which is known to have an i.v. half-time of only 8 minutes (*14*), within a lipid layer might improve local stability. In addition, we hypothesized that liposomal layers might function as mechanical entry barrier for SARS-CovV-2.

We therefore tested in a Syrian hamster SARS-Cov-2 infection model (*15*), whether NM +/- lipid carrier can lower the nasal virus load and subsequent COVID-19 severity.

## Material and Methods

### Animals

Hamsters have been previously described as model animals for COVID-19 (*15*). 8-9 weeks old female Syrian hamsters were purchased from Janvier Labs, France and included in the experiment after one week of acclimatization in the enhanced biosafety level 3 (BSL3) facility.

### Compounds

100mg Nafamostat mesylate (FUT-175, SelleckChem, Munich, Germany) was dissolved in 500 μl sterile water by gentle vortexing and subsequently diluted in 0.9% NaCl in water or lipid carrier (LipoNasal®, Optima Pharmazeutische GmbH, Hallbergmoos, Germany) to a final concentration of 1 mM.

### Virus

SARS-CoV-2 isolates (Human/NL/Lelystad/2020) were propagated in Vero E6 cells (ATCC) in Eagle’s minimal essential medium (Gibco) supplemented with 1% L-glutamine, 1% nonessential amino acids and 1% antibiotics, and 5% fetal bovine serum at 37 °C. All experiments with SARS-CoV-2 were performed in enhanced biosafety level 3 (BSL3) containment laboratories at the Wageningen Bioveterinary Research Facility, The Netherlands.

### Animal Treatment

The animal study was approved by the Dutch Central Authority for Scientific Procedures on Animals (CCD) prior to study start. Groups of each 10 female Syrian hamsters received under general anesthesia intranasal 100 μL NM (1mM) dissolved in 0.9% NaCL, 100 μL NM (1mM) dissolved in lipid carrier, 100 μl 0.9% NaCL, or 100 μl lipid carrier only.

5 minutes thereafter, all animals were infected by intranasal inoculation with 5 x 10^4^ TCID_50_ of SARS-COV-2. The animals were controlled daily for clinical signs of disease and weighed during 7 days. Hamsters were euthanized at more than 20% weight loss. On day 2 post infection (p.i.), two animals per group were sacrificed to collect respiratory tract tissues for histopathology. Nasal washes were collected on days 1 and 3 p.i. for viral load analysis.

### Virus titer analysis

Viral load in nasal washes was determined by RT-PCR. RNA was extracted from 500 uL of nasal wash fluid using the high Pure RNA Tissue Kit (Roche) and DNA was amplified in the Light Cycler (Roche) using SARS-CoV-2 Screening RTU kit (Kylt®). Viral RNA present in 10-fold dilution series of SARS-CoV-2 with known virus titer was analyzed in parallel to generate a standard curve for translation of cycle threshold (*C_τ_*) values into TCID_50_ titers.

### Histopathology and immunohistochemistry

Two animals from each group were necropsied on day 2 p.i. for histopathological examination. Formalin fixed nasal turbinates were embedded in paraffin, cut into 4 μm sections and stained with hematoxylin and eosin (HE).

For immunohistochemical staining of SARS-CoV2, sections were pretreated by heat-induced epitope retrieval in citrate buffer pH 6.0 (Vector Laboratories, Burlingame, USA) for 10 minutes at 121°C. Sections were then incubated with a rabbit polyclonal antibody directed against recombinant SARS-CoV nucleoprotein (40143-T62, Sino Biological, Beijing, China). A goat anti-rabbit HRP polymer (Envision +) was used as secondary antibody and color was developed with DAB + (DAKO / Agilent Technologies, Santa Clara, USA).

All slides were evaluated in a blinded fashion by a certified veterinary pathologist.

### Statistical analysis

Results represent mean ± standard deviation. All groups were compared using unpaired Student’s t-test (* p-value<0.05; ** p-value<0.01. Software used for calculation was GraphPad Prism version 8.

## Results

### Early nasal SARS-CoV-2 virus titers

Nasal washes were collected from each 10 animals per cohort at day 1 and each 8 animals per cohort at day 3 (2 were sacrificed for histopathology on day 2) and viral load measured by RT-PCR. At day 1 (Figure 1A) a significantly lower nasal virus load (as determined by log10/TCID50/ml) was detected in hamsters treated with NM in lipid carrier versus hamsters treated with saline solution. This difference was not detected at day 3 anymore (Figure 1B). Interestingly, numerical differences were also observed on day 1, when lipid carrier containing groups were compared with their NaCl control (lipid carrier versus NaCl, and NM in lipid carrier versus NM) (Figure 1A).

**Figure 1:**
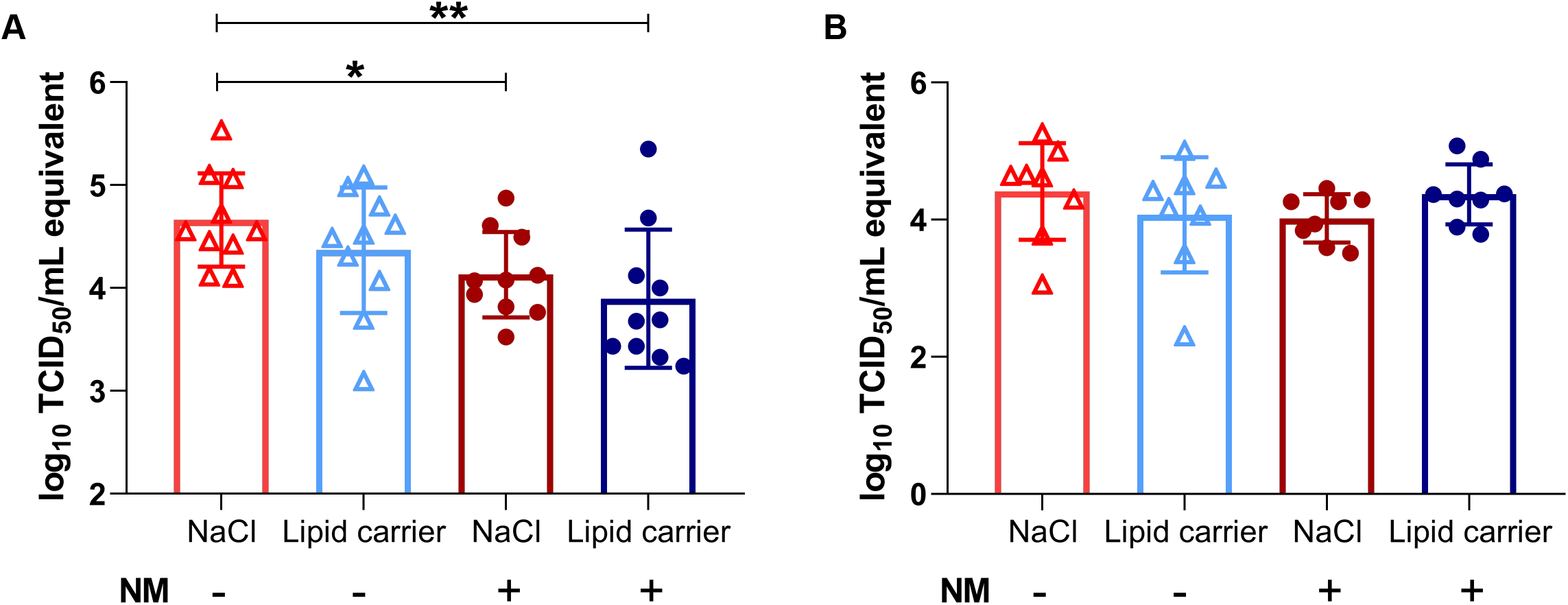
SARS-Cov2 nasal virus load is transiently reduced by NM and lipid carrier. The animals were analyzed by nasal wash for SARS-Cov2 nasal virus load on day 1 (n=10 per group) (A) and day 3 (n=8 per group) (B) after the indicated pre-treatment before nasal virus inoculation on day 0. Levels of viral RNA are expressed as log_10_ TCID_50_ per ml equivalents. Data shown as mean ± standard deviation. Unpaired t-test was used to compare across groups. * p-value= 0.0135, **p-value= 0.0078. NM Nafamostat mesylate

### Body weight

Weight loss has been described previously to be associated with severity and disease progression in the SARS-CoV2 hamster model (*15*). The average weight in each cohort was similar pre-infection. Post infection all animals lost weight until day 7 (independent of the treatment with NM or lipid or NaCl) or had to be sacrificed before day 7 due to achieving the maximum allowed weight loss of 20% (Figure 2).

**Figure 2:**
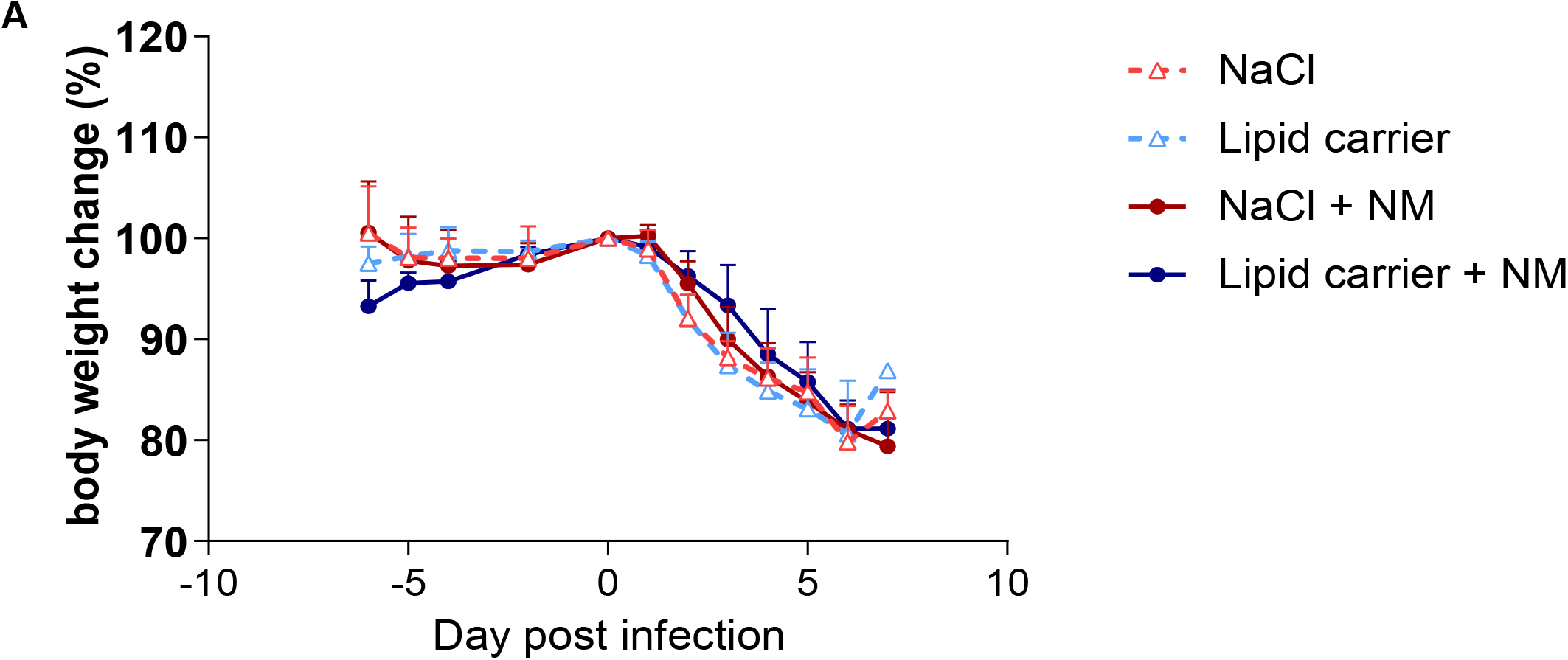
Body weight loss is not prevented by single Nafamostat mesylate +/- lipid carrier nasal application. Each 10 animals (8 animals from day 2 on) were pretreated by single nasal treatment application as indicated and daily weighed post nasal SARS-CoV2 virus inoculation. All animals were euthanized by day 7 p.i. or when they reached clinical endpoint (>20% weight loss relative to day 0). Error bars represent standard deviation. NM Nafamostat mesylate.

### Histopathology

Histopathology of the nasal cavity in hamsters (n = 2) 2 days p.i. showed no differences between the groups. Representative pictures of the catarrhal rhinitis in all groups with edema, in some cases fibrin in the lamina propria under the nasal epithelium, and transmigration of neutrophilic granulocytes from the blood vessels into and across the nasal epithelial lining are shown in Figure 3A and 3B. Also, local loss of integrity of the nasal epithelium with degeneration/ necrosis of epithelial cells was observed (Figure 3C). Abundant exudate was regularly present within the nasal cavity, containing mainly neutrophils and some macrophages.

**Figure 3.**
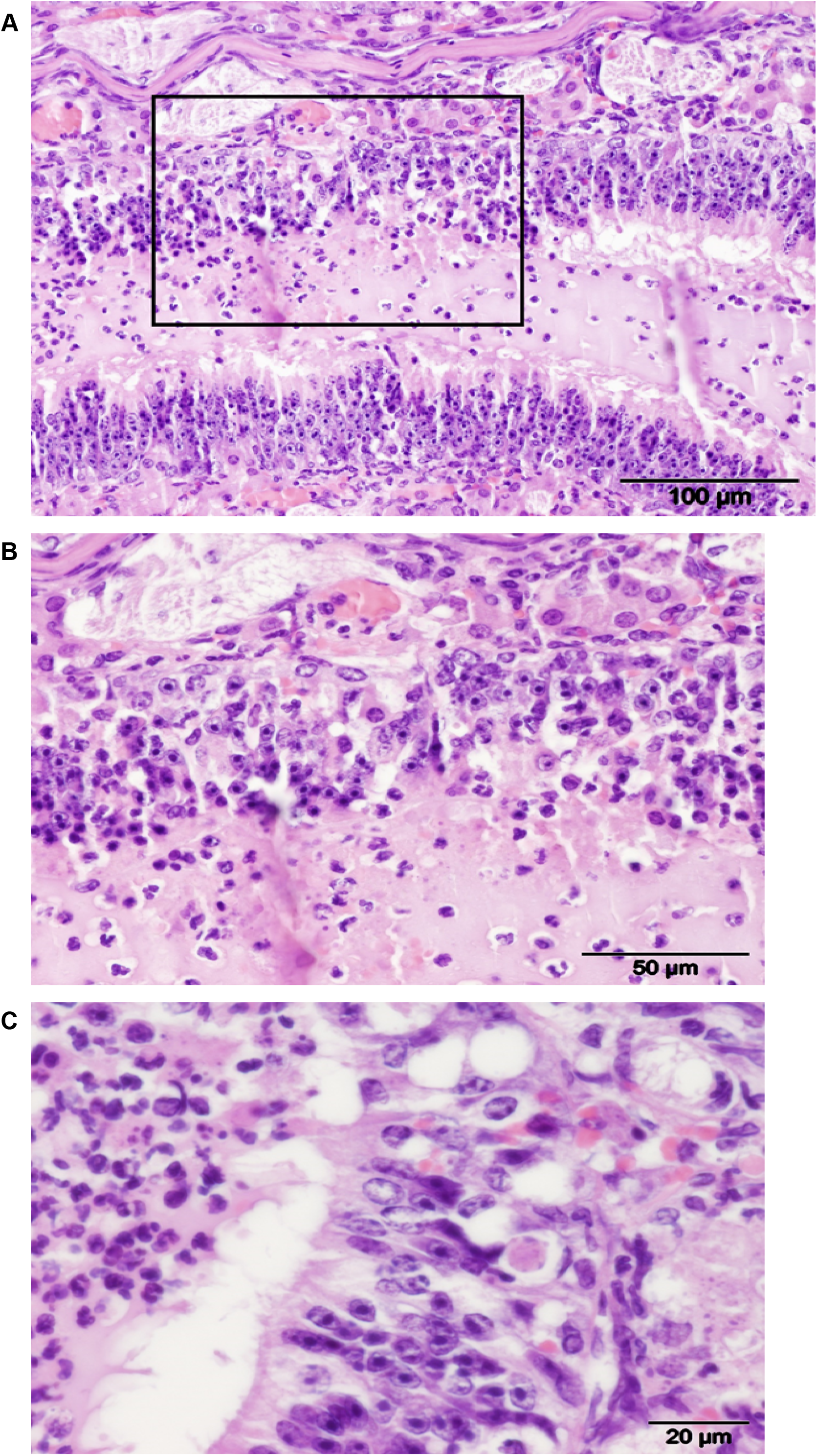
Histopathology analysis of nose caveat on day 2. (A) Acute catarrhal rhinitis. Exudative inflammation of nasal mucosa. Note the exudate with granulocytes and macrophages in the nasal cavity lumen.(B) Magnification of area in the rectangle of Figure 3A. Transmigration of neutrophilic granulocytes from the blood vessels of the lamina propria into and across the nasal epithelium. (C) Local degeneration and necrosis of nasal epithelial cells.

### Immunohistochemistry for SARS-Cov2 nucleoprotein

Independent of pre-treatment, in all four groups almost the entire nasal epithelium stained strongly positive for the SARS-CoV2 nucleoprotein (75-100% in each animal as determined in a blinded fashion). This involved both the pseudostratified respiratory epithelium of the nose and sinuses, as well as the predominantly present pseudostratified olfactory epithelium (Figure 4 A-C).

**Figure 4.**
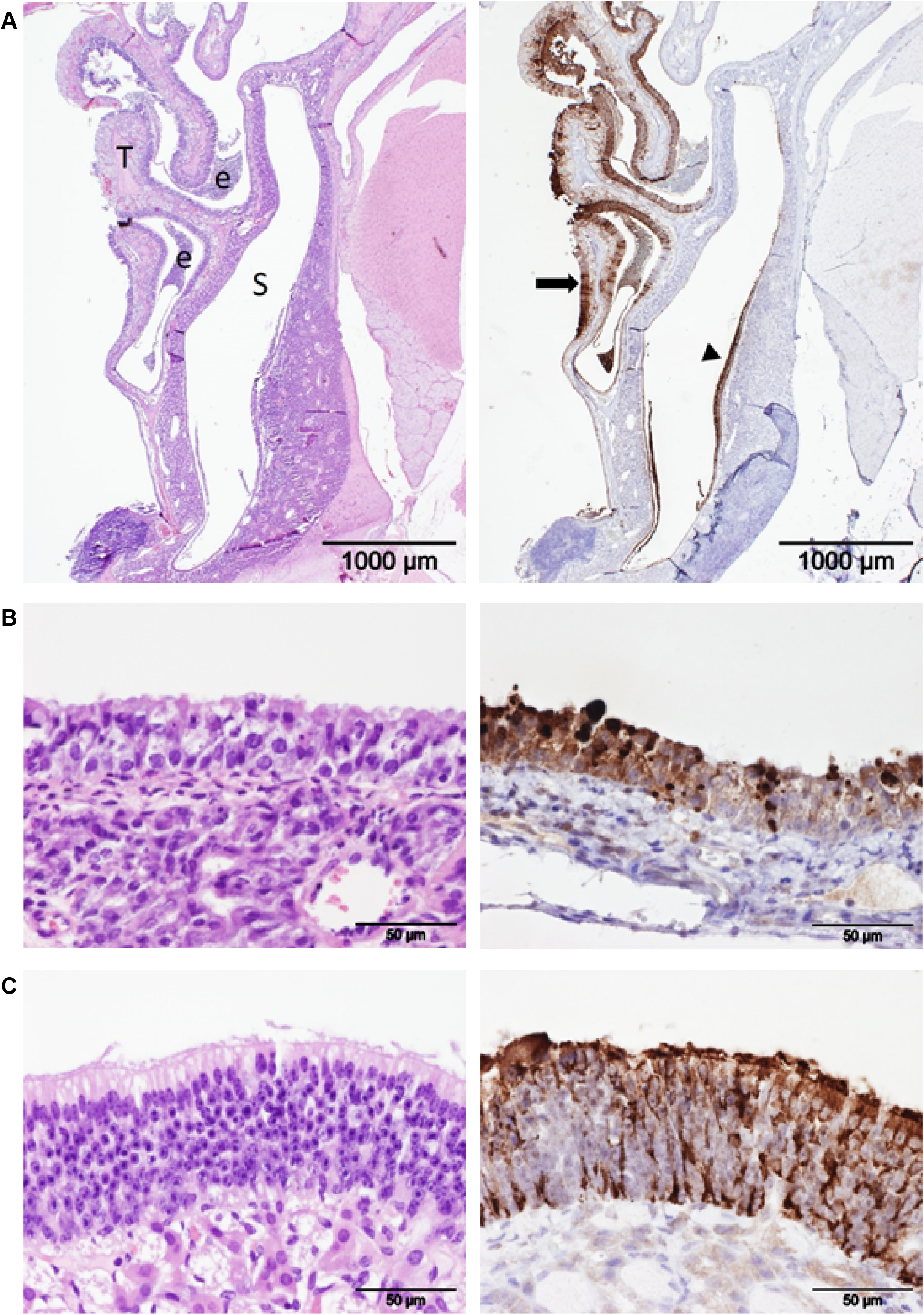
Immunohistologic staining for SARS-CoV2. (A) Serial section of the nose at low magnification stained with HE (left) or immunostained for SARS-CoV2 (right). Note the strong staining of the olfactory (arrow) and respiratory epithelium (arrow head) for the SARS-CoV2 nucleoprotein. S = maxilllary sinus; T = nasal turbinates, e = exudate within the nasal cavity. (B) Nasal respiratory epithelium stained with HE (left) and immunostained for SARS-CoV2 antigen. (C) Nasal olfactory epithelium stained with HE (left) and immunostained for SARS-CoV2 antigen

## Discussion

The SARS-CoV2 pandemic has not only become a threat to the health of mainly older people all over the world, but has also transformed strongly current open societies requiring social distancing, home office working, wearing face masks, and full lock-downs. This is in strong contrast to the peoples’ wishes of gathering together and interact socially, as resembled/ reflected by the decreasing acceptance in daily-life restrictions requested by the governments during this second wave of the pandemic, and increasing frequencies of demonstrations against them.

Alternative approaches, like chemical virus entry restrictions/blocking have several advantages; first, they could allow more social interaction, second, they could form a bridge until effective vaccines are developed, and third, they could even be used in combination with vaccination or local antibody delivery. Furthermore, blocking viral entry stops infection early on, preventing viral replication. As many viruses exploit cellular endocytic mechanisms to initiate internalization and infection, and cells have just a few such mechanisms, inhibiting these pathways may even affect different viruses (*16*).

Nafamostat mesylate (NM) has been shown to effectively inhibit SARS-CoV-2 cell entry in in *vitro* experiments (*9*). This was achieved with NM levels of above 0.1-1 μM. Considering the fact, that the half-life of NM is i.v. only 8 minutes (*8*), a consistent effect might be only achieved by repetitive nasal application of NM 1 mM every hour, to achieve local levels resembling the *in vitro* experiments. Therefore, we tested in our experiment the local nasal application of NM at a 1 mM dosing.

While we observed a significant reduction of the viral load in the nasal cavity, this reduction was only transient, and did not significantly reduce the onset/degree/severity of rhinitis and weight loss in our experimental setting. Also, the addition of a lipid carrier did not significantly reduce the virus titers. There might be several reasons for the incongruence of virus load reduction and clinical benefit. First, the reduction was not strong enough, to achieve clinical effects, second, a single application is not sufficient (unfortunately repetitive anesthesia that is required for the nasal application is not allowed by law in the Netherlands) or third, the hamster experiment simulated not an early nasal infection, but late spreading due to the fact that the droplet with the virus directly moves through the throat to the lungs. A transmission experiment, as just recently described (*17*), might better reflect the situation in practice in terms of dose and route. On the other hand, timing of the NM treatment to infection interval is less controllable. Unfortunately, this was not known at the time of the design of the experiment reported here, and we had no access to a ferret transmission experiment. Indeed, a transmission experiments in ferrets testing the nasal application of a lipopeptide was significantly positive (de Vries et al., bioRxiv preprint doi: https://doi.org/10.1101/2020.11.04.361154). Thus, NM in combination with such a lipopeptide approach might be even more effective.

Despite the current lack of *in vivo* data, and in line with our idea, the university of Tokyo partnered with several pharmaceutical companies to develop an inhalative formulation of NM (https://www.u-tokyo.ac.ip/focus/en/articles/zl30400140.html).

Several other approaches of local therapy for prevention of COVID-19 are currently underway. Probably most promising is local application of an anti-protein S nanobody for inhalation (*12*), or the above discussed inhibitory lipopeptide (de Vries et al.). A major advantage of such different approaches is, that they even could be even combined, as long as they mediate no local immune suppression.

In summary, our experiments for the first time indicate *in vivo* that NM in saline solution or in a lipid carrier applied as nose spray or inhalator can reduce the nasal viral load, which might reduce the risk for transmission, nasal viral load/ infection and severity of COVID-19 after exposure to SARS-CoV-2. However, optimization of local stability will be a prerequisite for clinical success.

## References

1. W. Li et al., Angiotensin-converting enzyme 2 is a functional receptor for the SARS coronavirus. Nature 426, 450–454 (2003).

2. M. Hoffmann et al., SARS-CoV-2 Cell Entry Depends on ACE2 and TMPRSS2 and Is Blocked by a Clinically Proven Protease Inhibitor. Cell 181, 271–280 e278 (2020).

3. Y. J. Hou et al., SARS-CoV-2 Reverse Genetics Reveals a Variable Infection Gradient in the Respiratory Tract. Cell 182, 429–446 e414 (2020).

4. R. L. Tillett et al., Genomic evidence for reinfection with SARS-CoV-2: a case study. Lancet Infect Dis, (2020).

5. M. Mulder et al., Reinfection of SARS-CoV-2 in an immunocompromised patient: a case report. Clin Infect Dis, (2020).

6. K. K. To et al., COVID-19 re-infection by a phylogenetically distinct SARS-coronavirus-2 strain confirmed by whole genome sequencing. Clin Infect Dis, (2020).

7. C. Xia, S. S. Lam, C. Sonne, Ban unsustainable mink production. Science 370, 539 (2020).

8. K. Okajima, M. Uchibat, K. Murakamit, Nafamostat Mesilate. Cardiovascular Drug Reviews 13, 51–65 (1995).

9. M. Yamamoto et al., Identification of Nafamostat as a Potent Inhibitor of Middle East Respiratory Syndrome Coronavirus S Protein-Mediated Membrane Fusion Using the Split-Protein-Based Cell-Cell Fusion Assay. Antimicrobial agents and chemotherapy 60, 6532–6539 (2016).

10. M. Hoffmann et al., Nafamostat Mesylate Blocks Activation of SARS-CoV-2: New Treatment Option for COVID-19. Antimicrobial agents and chemotherapy 64, (2020).

11. M. Yamamoto et al., The Anticoagulant Nafamostat Potently Inhibits SARS-CoV-2 S Protein-Mediated Fusion in a Cell Fusion Assay System and Viral Infection In Vitro in a Cell-Type-Dependent Manner. Viruses 12, (2020).

12. M. Schoof et al., An ultra-high affinity synthetic nanobody blocks SARS-CoV-2 infection by locking Spike into an inactive conformation. bioRxiv, (2020).

13. M. Bohm, G. Avgitidou, E. El Hassan, R. Mosges, Liposomes: a new non-pharmacological therapy concept for seasonal-allergic-rhinoconjunctivitis. European archives of oto-rhino-laryngology: official Journal of the European Federation of Oto-Rhino-Laryngological Societies 269, 495–502 (2012).

14. A. T., Nafamostat mesilate. Drugs of the Future 9, 747–748 (1984).

15. M. Imai et al., Syrian hamsters as a small animal model for SARS-CoV-2 infection and countermeasure development. Proc Natl Acad Sci U S A 117, 16587–16595 (2020).

16. M. Mazzon, M. Marsh, Targeting viral entry as a strategy for broad-spectrum antivirals. F1000Research 8, (2019).

17. J. F. Chan et al., Simulation of the clinical and pathological manifestations of Coronavirus Disease 2019 (COVID-19) in golden Syrian hamster model: implications for disease pathogenesis and transmissibility. Clin Infect Dis, (2020).

